# Visually Induced Changes in Cytokine Production in the Chick Choroid

**DOI:** 10.1101/2021.06.03.446867

**Authors:** Jody A. Summers, Elizabeth Martinez Cano

## Abstract

Postnatal ocular growth is regulated by a vision-dependent mechanism which acts to minimize refractive error through coordinated growth of the ocular tissues. Of great interest is the identification of the chemical signals that control visually-guided ocular growth. Here we provide evidence that the pro-inflammatory cytokine, Interleukin-6 (IL-6), may play a pivotal role in the control of ocular growth using a chicken model of myopia. Microarray, real time RT-qPCR, and ELISA analyses identified IL-6 upregulation in the choroids of chick eyes under two visual conditions that introduce myopic defocus and slow the rate of ocular elongation (recovery from induced myopia and compensation for positive lenses). Intraocular administration of atropine, an agent known to slow ocular elongation, also resulted in an increase in choroidal IL-6 gene expression. Nitric oxide appears to directly or indirectly upregulate choroidal IL-6 gene expression, as administration of the non-specific nitric oxide synthase inhibitor, L-NAME, inhibited choroidal IL-6 gene expression, and application of a nitric oxide donor stimulated IL-6 gene and protein expression in isolated chick choroids. Considering the pleiotropic nature of IL-6 and involvement in many biological processes, these results suggest that IL-6 may mediate many aspects of the choroidal response in the control of ocular growth.

## Introduction

High myopia is a significant risk factor for several blinding eye diseases including glaucoma, retinal detachment and macular degeneration, and therefore represents a leading cause of blindness worldwide (Buch, Vinding et al., 2001). The prevalence of myopia is continuing to increase and is expected to affect nearly half of the global population by 2050 (Holden, Fricke et al., 2016). Although clinical and experimental studies indicate that normal eye growth (emmetropization) is controlled by visual input (Wallman & Winawer, 2004), the cause of myopia in humans is not understood.

Animal models have provided valuable insights into the role of the visual environment on ocular growth control. Deprivation of form vision, through the use of visual “occluders” or “goggles” results in accelerated ocular growth and the development of myopia within a matter of days in chicks, tree shrews, guinea pigs, and primates (Howlett & McFadden, 2006, Sherman, Norton et al., 1977, Troilo & Judge, 1993, Wallman, Turkel et al., 1978). Upon removal of the occluder, normal visual input is restored to the elongated myopic eye, resulting in a rapid deceleration in ocular elongation and eventual return to emmetropia (“recovery”) (Wallman & Adams, 1987).

Postnatal ocular growth can also be manipulated through the application of positive or negative lenses, as the eye has been shown to compensate for the imposed defocus in many vertebrates, including fish, chicks, mammals, and primates (Graham & Judge, 1999, Hung, Crawford et al., 1995, Norton, Siegwart et al., 2006, Schaeffel, Glasser et al., 1988, Shen & Sivak, 2007). Application of positive lenses, which cause images to form in front of the retina (myopic defocus), results in slowing the rate of ocular elongation and thickening of the choroid, in order to push the retina toward the image plane (Wallman, Wildsoet et al., 1995). Conversely, application of negative lenses results in an increased rate of elongation and thinning the choroid to pull the retina back toward the image plane.

It is well-established that visually induced changes in ocular length are the result of a locally driven “retina-to-choroid-to-scleral chemical signaling cascade” that is initiated by a visual stimulus, followed by chemical changes in the retina and choroid, ultimately resulting in altered extracellular matrix (ECM) remodeling of the scleral shell (Troilo, Smith et al., 2019). The choroid, a highly vascularized layer located immediately adjacent to the sclera, has been shown to undergo changes in thickness, permeability and blood flow during periods of visually guided eye growth (Fitzgerald, Wildsoet et al., 2002, Pendrak, Papastergiou et al., 2000, Rada & Palmer, 2007, Wallman et al., 1995). Moreover, due its proximity to the sclera, the choroid is suspected to synthesize and/or release scleral growth regulators to control the rate of ocular elongation in response to visual stimuli (Marzani & Wallman, 1997, Rada & Palmer, 2007). All-*trans*-retinoic acid is one potential choroidally-derived scleral growth regulator, whose choroidal concentrations are modulated by the activity of retinaldehyde dehydrogenase 2 (RALDH2)(Mertz & Wallman, 2000, Rada, 2012). Of much interest, therefore, are the identification of genes causally involved in the regulation of the choroidal response during visually guided eye growth.

Here, we report rapid and significant changes in choroidal gene expression of the pluripotent cytokine, interleukin 6 (IL-6), in response to myopic defocus and in response to chemical treatments known to modulate eye growth. Considering the pleiotropic nature of IL-6 and involvement in many biological processes, these results suggest that IL-6 may mediate many aspects of the choroidal response in the control of ocular growth. A preliminary report of our findings was presented previously (Summers and Martinez, IOVS 2020 61: E-abstract 3400).

## Results

### Expression of IL-6 in chick ocular tissues

Immunohistochemical staining for IL-6 indicated that IL-6 is expressed in numerous cells throughout the choroid and RPE as punctate cytoplasmic deposits (**Figure 1A,B**). IL-6-containing cells included the RPE, choroidal endothelial cells (blood vessels are labelled with asterisks), and choroidal stromal cells. Immunolabelling was abolished after incubation of this antibody with a 10-fold molar excess of chicken IL-6 demonstrating that the immunohistochemical detection procedure was specific (**Figure 1C**).

**Figure 1.**
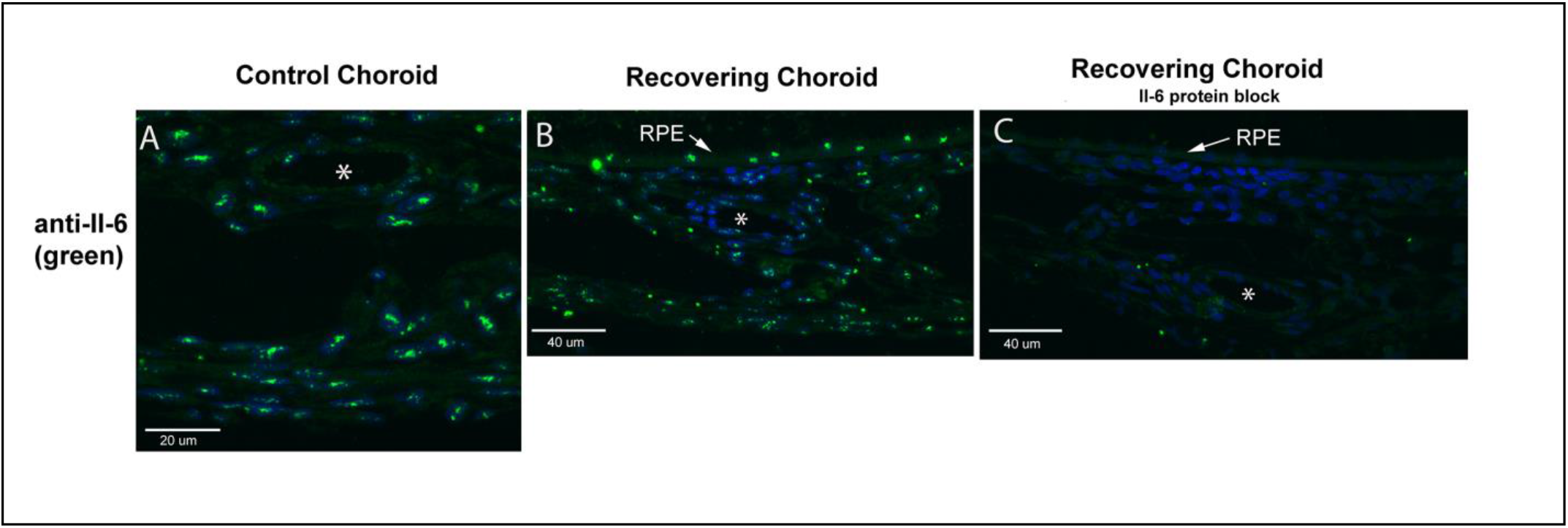
Immunohistochemical localization of IL-6 in chick choroids. (**A, B**) Il-6 was localized in treated and control eyes after 1 day of recovery from induced myopia. (**C**) Preincubation of anti-IL-6 with a 10 fold molar excess of recombinant chicken IL-6 (1.67 μM) before use on tissue sections abolished IL-6 labeling. Bar = 20 μm in **A** and 40 μm in **B, C**. Choroidal blood vessels are indicated by asterisks (*). RPE = retinal pigmented epithelium.

### IL-6 is up-regulated following visual stimuli

#### Form Deprivation and Recovery

Preliminary results from an Affymetrix microarray experiment indicated that IL-6 was increased over 10 fold in choroids of chick eyes following 6 hrs of recovery, compared with normal, untreated eyes (**Figure 2**).

**Figure 2.**
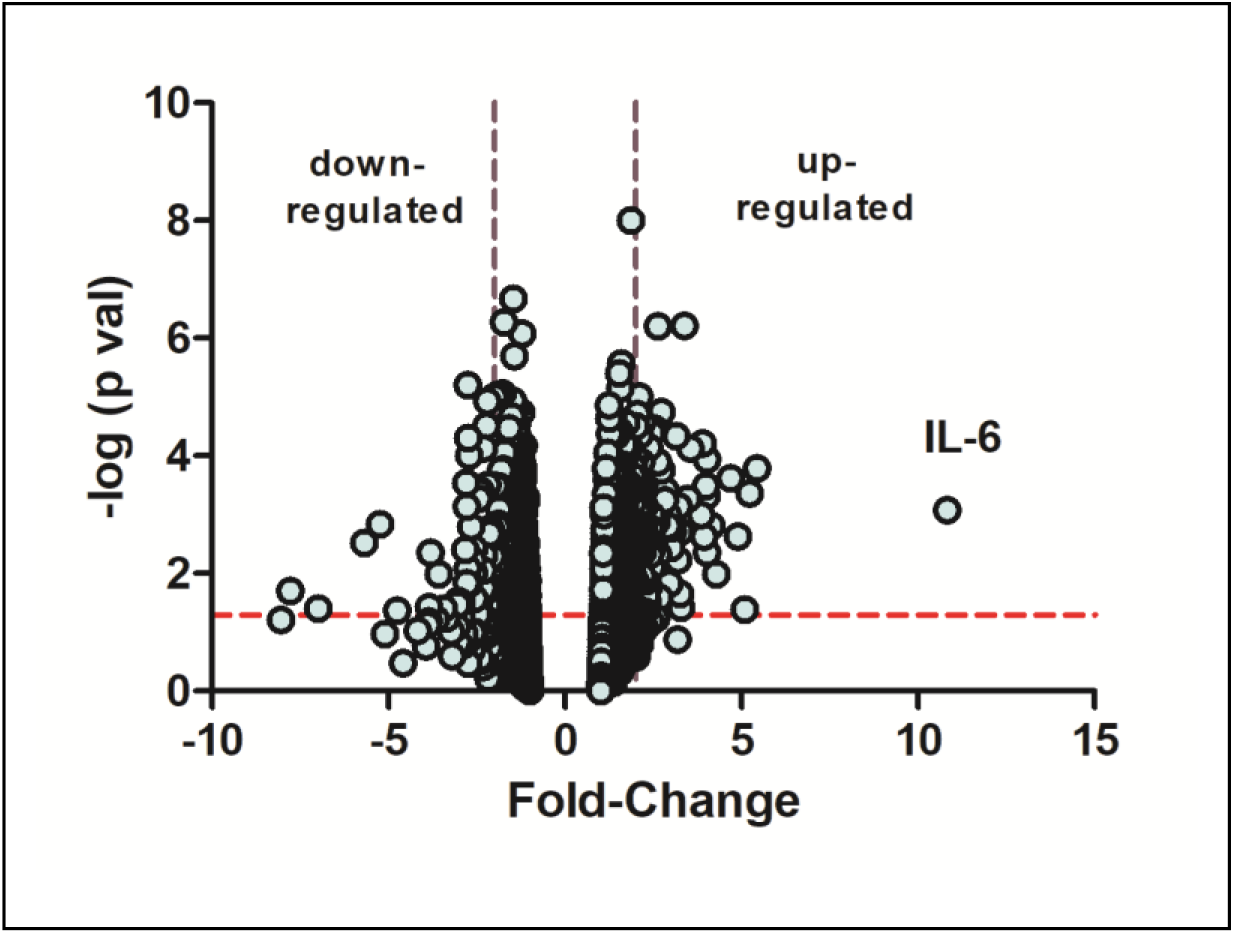
Microarray identifies IL-6 as a gene highly overexpressed in early recovery. A volcano plot of Affymetrix chicken microarray data indicated that 207 genes were found to be significantly differentially expressed by ≥ 2 fold in recovering choroids as compared with choroids from normal untreated chicks (p ≤ 0.05). The horizontal dashed red line indicates where p = 0.05, with points above the line having p < 0.05 and points below the line having p > 0.05. The area between the dashed purple lines indicates points having a fold-change less than |2|. IL-6 was increased by 10.83 fold in recovering choroids compared with normal choroids (n = 5 birds in each group) p = 0.00084, Student’s t-test).

To determine the precise temporal pattern of IL-6 expression during recovery, we utilized Taqman™ real time PCR to quantify IL-6 mRNA concentrations in choroids following 10 days of form deprivation and over several time points during recovery from induced myopia (**Figure 3A**). IL-6 mRNA was significantly increased in choroids following 45 minutes to 24 hrs of recovery compared to contralateral control eyes (↑99 - 1738%), reaching a maximum following 6 hrs of recovery. By 4 days of recovery, IL-6 mRNA was significantly downregulated in treated choroids, compared with that of treated choroids at 24 hrs of recovery, and was similar to that of fellow control eyes (p = 0.84 and 0.12, for 4 and 8 days of recovery, respectively, paired t-test).

**Figure 3.**
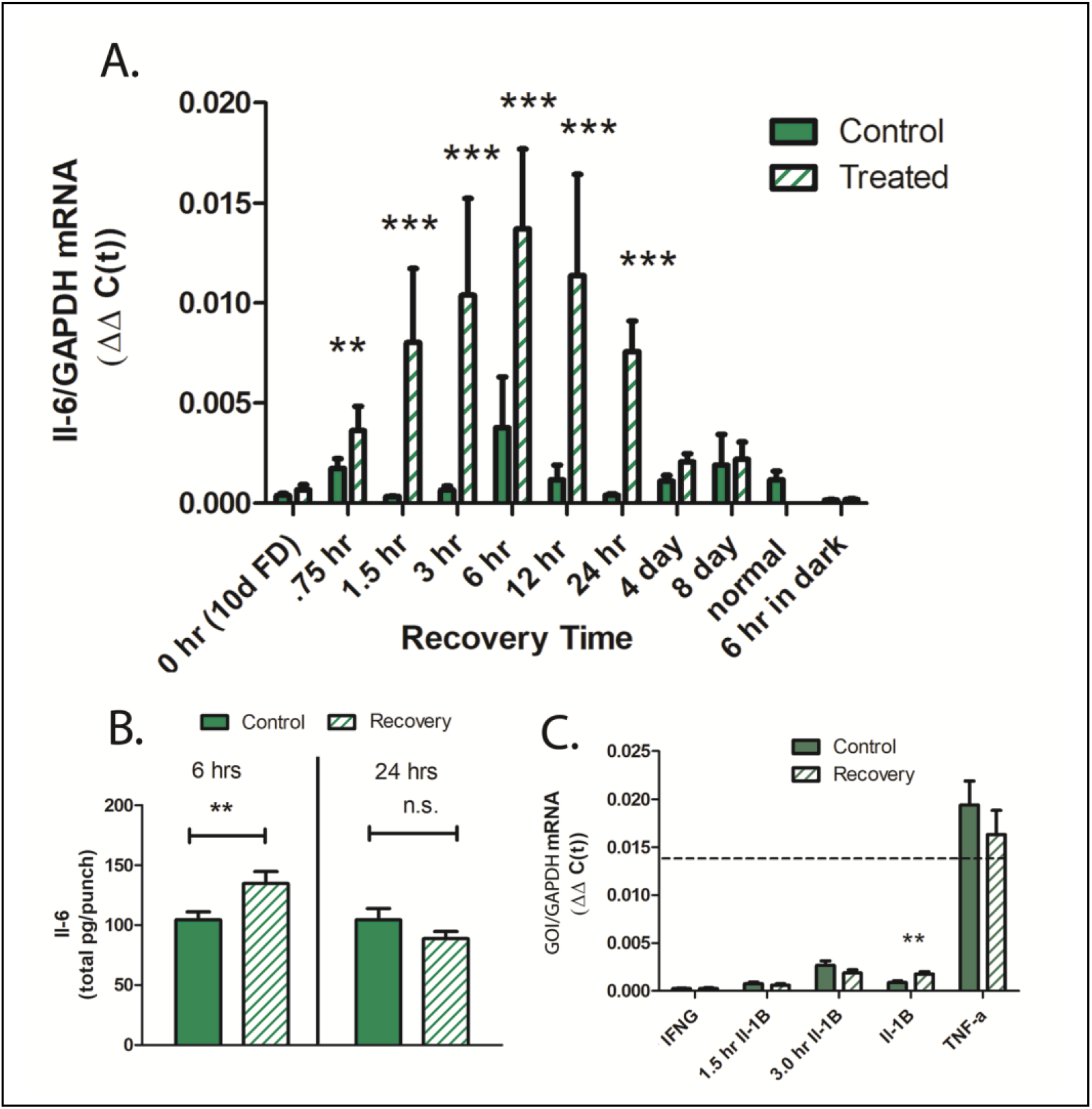
Cytokine gene and protein expression in chick choroids. (A) IL-6 mRNA expression in choroids from control and treated eyes, following 10 days of form deprivation (0 hr/10d FD), 0.75 hr – 8 days of recovery from form deprivation, normal, untreated eyes (normal), and in eyes recovered for 6 hrs, but kept in total darkness (6 hr in dark) (n = 5 – 16 birds in each group) *** p < 0.001, ** p < 0.01, paired t-test. (B) IL-6 protein production by control and recovering choroids following 6 and 24 hrs of recovery from induced myopia. Data are expressed as mean ± SEM (n = 16) ** p < 0.01, paired t-test. (C) Quantification of other proinflammatory cytokines in chick choroids. Gene expression of Interferon gamma (IFNG), interleukin 1B (IL-1B), and tumor necrosis factor alpha (TNF-a) was quantified in control and treated chick choroids following 6 hrs of recovery. Additionally, IL-1B mRNA was quantified following 1.5 and 3 hr of recovery. The dashed line indicates the average IL-6 expression in 6 hr recovering choroids. (n = 6 – 11 birds in each group) ** p < 0.01, paired t-test.

The rapid increase in choroidal IL-6 gene expression observed during recovery, i.e. 0.75 hrs following removal of the occluder, prompted us to determine whether increased choroidal IL-6 gene expression was an artifact of removal of the occluder, rather than due to a visual stimulus. To address this possibility, one group of chicks was kept in complete darkness for 6 hrs following removal of the occluder (**Figure 3A,** “6 hrs in dark”). Interestingly, IL-6 gene expression was significantly lower in both control and recovering eyes, as compared with IL-6 mRNA levels from all other choroids of control, recovering or form deprived eyes reared under normal room light [p < 0.01, t-test, for choroids of dark reared control or recovering eyes compared with the lowest control group (1.5 hr control group)].

Choroidal protein expression of IL-6 was also significantly increased following 6 hrs of recovery, compared to contralateral control eyes (↑ 36 %, p < 0.01, paired t-test), but returned to control levels by 24 hrs of recovery (**Figure 3B**). We also evaluated gene expression of the chicken cytokines, interferon gamma (IFN-γ), interleukin 1β (IL-1 β), and TNF-α (TNF-α) in choroids of eyes following 1.5 – 6 hrs of recovery and in contralateral control eyes. Gene expression of TNF-α was substantially higher (≈ 7 fold) than all other cytokines examined, but not significantly different between control and recovering eyes. Only gene expression of IL-1 β following 6 hours of recovery was significantly elevated in recovering eyes compared with controls (↑ 179 %, p<0.05, paired t-test) **(Figure 3C**).

#### Light Intensity

Based on our observation that IL-6 mRNA was significantly lower in choroids of birds kept in darkness for 6 hrs, compared to control or treated eyes reared under standard room lighting, we evaluated the effect of varied light intensity on choroidal IL-6 gene expression. Normal untreated chicks were kept in dim light (5 lux), medium intensity light (700 lux) and high intensity light (3150 lux), as well as red LED light (58 lux) and blue LED light (111 lux) for 6 hours prior to RNA isolation. Exposure to all light intensities resulted in a significant increase in IL-6 mRNA, compared to IL-6 gene expression in choroids of dark reared chicks (**Figure 4**); however, no differences were observed in IL-6 mRNA levels between the five lighting conditions, with all IL-6 mRNA values similar to that of the normal untreated chick choroids (**Figure 3A**).

**Figure 4.**
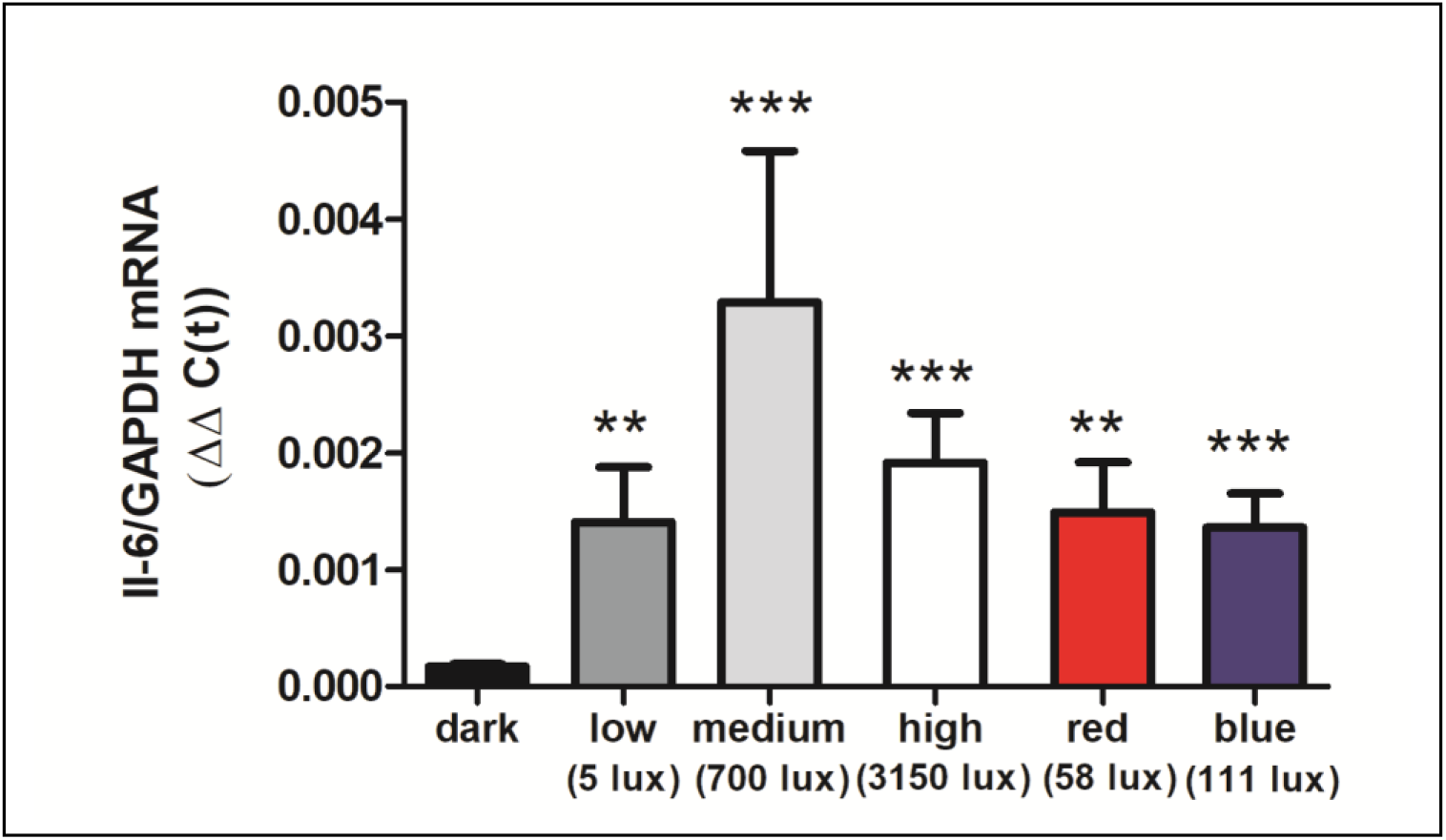
Effect of light intensity on IL-6 mRNA expression. Normal chicks were housed in complete darkness (“dark”), white LED light of varying intensities (“low”, 5 lux; “medium”, 700 lux; “high”, 3150 lux), red LED light (“red”, 58 lux), or blue LED light (“blue”, 111 lux) for 6 hrs at which time choroids were isolated with Il-6 mRNA was quantified by Taqman™ real time PCR. [n = 6 – 8 birds (12 – 16 choroids)] in each group. *** p < 0.001, ** p < 0.01, Kruskal-Wallis test with Dunn’s multiple comparisons.

#### Optical Defocus

Following removal of the occluder, previously form deprived eyes experience myopic defocus due to form deprivation-induced myopia. We therefore determined whether choroidal IL-6 gene expression was affected following a period of imposed myopic or hyperopic defocus via the application of +15D or −15D spectacle lenses (**Figures 5A & 5B**). Following 24 hrs of +15D lens wear, choroidal IL-6 gene expression was significantly increased compared with contralateral control eyes (↑48.3%, p<0.05, paired t-test) **(Figure 5C).** No significant differences were detected in IL-6 gene expression following 6 hrs of +15D lens wear. Treatment with −15D lenses had no statistically significant effect on choroidal IL-6 gene expression, although a trend toward decreased expression was noted. Scleral proteoglycan synthesis was also assessed following 24 hrs of lens treatment to confirm that the +15D and −15D lenses were inducing compensatory ocular growth responses **(Figure 5D).** As expected, treatment with +15D lenses resulted in a significant decrease in scleral proteoglycan synthesis (P<0.001, paired t-test) and treatment with −15D lenses resulted in a significant increase in scleral proteoglycan synthesis (P<0.05, paired t-test).

**Figure 5.**
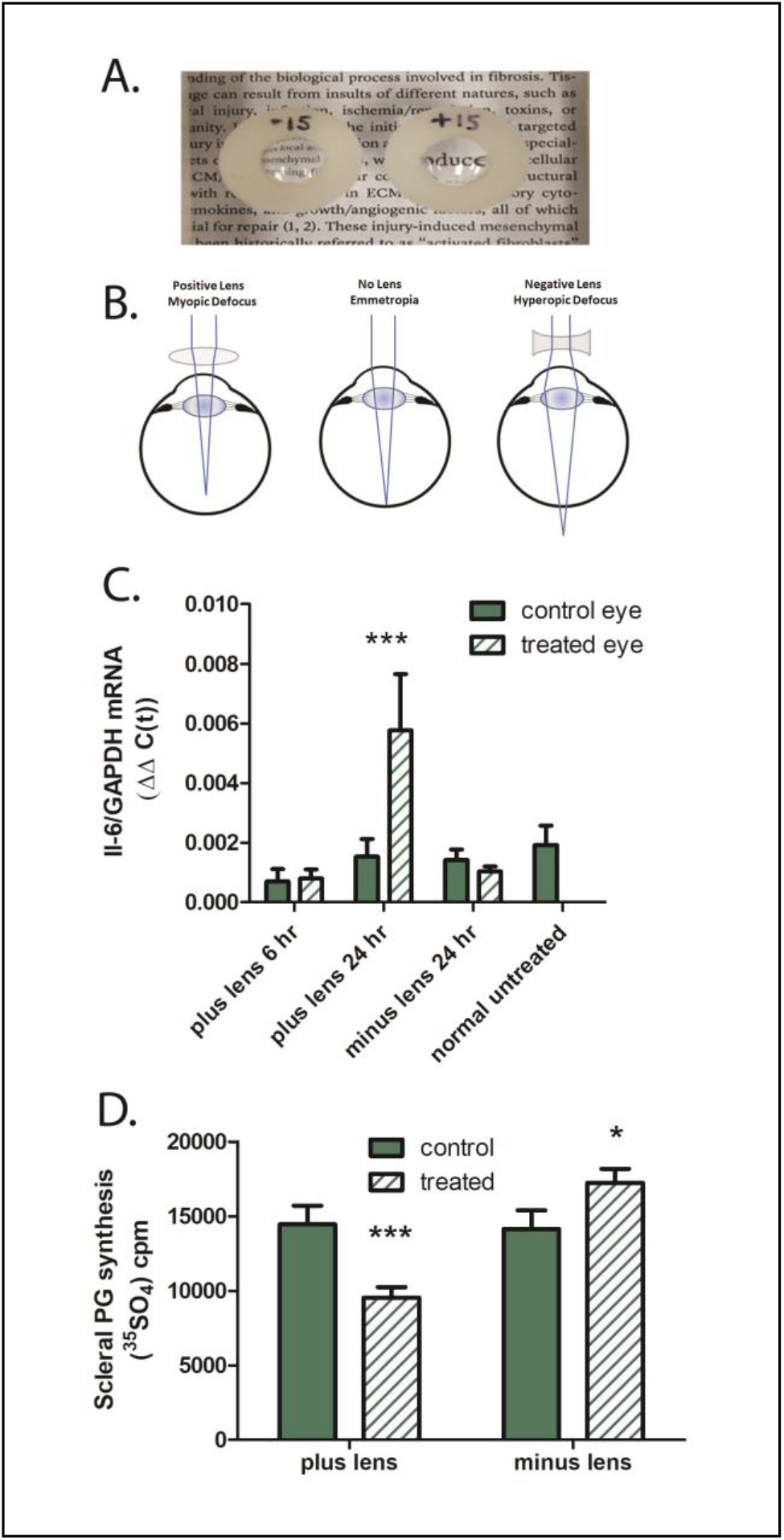
Effect of imposed defocus on choroidal IL-6 gene expression. **(A).** Spectacle lenses [minus 15D (−15) or plus 15D (+15)] were applied to the right eyes of chicks for 6 – 24 hrs. **(B).** Schematic diagram illustrating the effects of imposed optical defocus on the location of ocular images of distant objects for an emmetropic eye (center); positive lenses move the image plane in front the retina, imposing myopic defocus (left), while negative lenses move the image plane behind the retina, imposing hyperopic defocus (right). **(C)** IL-6 mRNA expression in choroids from control and treated eyes, following 6 or 24 hr of plus lens wear (n = 6 and n = 27, respectively), 24 hr of minus lens wear (n = 34), and normal untreated choroids. (n = 8). *** p < 0.001, paired t-test. **(D)** Scleral proteoglycan synthesis following 24 hrs of lens wear. Proteoglycan synthesis was significantly reduced following 24 hrs of +15D lens wear, compared to untreated contralateral control eyes (***p < 0.001, paired t-test, n = 10) and was significantly increased following 24 hrs of −15D lens wear, compared with untreated contralateral control eyes (* p < 0.05, paired t-test, n = 13).

### Choroidal IL-6 Synthesis Is Transcriptionally Regulated by NO

Nickla et al., (Nickla, Damyanova et al., 2009) have previously demonstrated that nitric oxide synthesis via neuronal nitric oxide synthase (nNOS) is obligatory for recovery from form deprivation myopia. Administration of the non-specific inhibitor nitric oxide synthase inhibitor, N^a^-nitro-L-arginine methyl ester (L-NAME), or the nNOS inhibitor N^w^ -propyl-L-arginine, blocks recovery due to inhibition of choroidal thickening and dis-inhibition of scleral proteoglycan synthesis. We therefore investigated the role of nitric oxide on choroidal IL-6 transcription using several approaches. First, L-NAME, or vehicle, was administered via intravitreal injection to chick eyes following 10 days of form deprivation. Chicks were then given unrestricted vision for 6 hours and choroidal IL-6 mRNA was quantified **(Figure 6A).** Following 6 hrs of recovery, IL-6 mRNA was significantly increased in recovering eyes of vehicle (saline) treated eyes compared with contralateral control eyes (↑1107%, p<0.0001, paired t-test). Administration of L-NAME just prior to recovery resulted in a significant decrease in choroidal IL-6 mRNA, 6 hrs following L-NAME administration, as compared with choroidal IL-6 mRNA levels in recovering eyes of saline-treated eyes (p <0.05, Mann Whitney U test). L-NAME administration did not completely abolish the recovery-induced rise in choroidal IL-6 mRNA; IL-6 mRNA levels in choroids of L-NAME-treated eyes were significantly higher than that of contralateral untreated eyes (p < 0.01, paired t-test). As previously reported (Summers Rada & Hollaway, 2011), scleral proteoglycan synthesis was significantly increased in the posterior sclera of chick eyes during the development of form deprivation myopia (Day 0 of recovery) (p < 0.001, paired t-test) and was rapidly downregulated following 12 hrs of recovery to levels similar to that of contralateral control eyes (vehicle) (**Figure 6B**). Intravitreal application of L-NAME inhibited this recovery response, resulting in a significant increase in scleral proteoglycan synthesis in recovering eyes, as compared with contralateral control eyes (p<0.01, paired t-test) and compared with recovering eyes of vehicle-treated chicks (p < 0.05, Mann-Whitney U test). These results confirm that intravitreal administration of L-NAME in our study resulted in the same effects on eye growth as have been previously reported.

**Figure 6.**
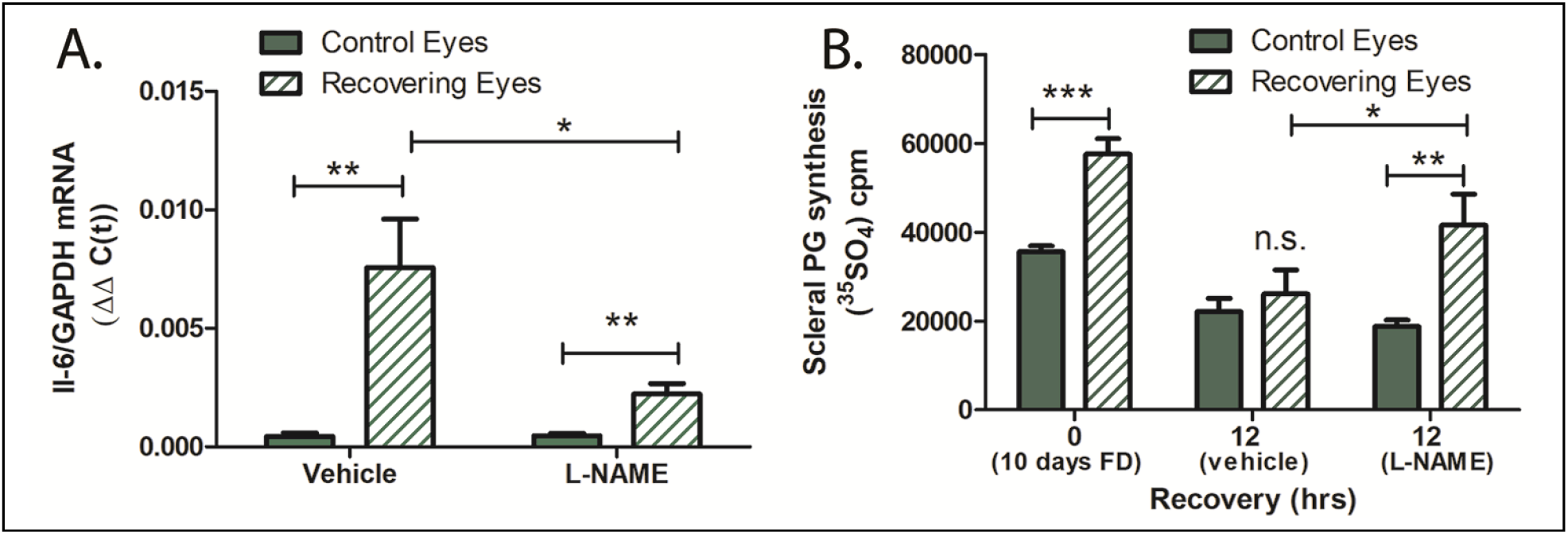
L-NAME inhibits choroidal IL-6 transcription and recovery. **(A)** Intravitreal injection of L-NAME (16.2 μmole/eye) immediately prior to recovery significantly reduced IL-6 mRNA levels compared to recovering eyes receiving vehicle only (0.9% NaCl) (*p < 0.05, Mann-Whitney U test, n = 7; **p < 0.01, paired t-test, n = 7). **(B)** L-NAME disinhibits scleral proteoglycan synthesis in recovering eyes. Following 12 hrs of recovery from 10 days of form deprivation (FD), scleral proteoglycan synthesis decreased to control levels in vehicle-treated eyes, but remains significantly increased over control levels in L-NAME treated eyes. (*** p < 0.001, ** p< 0.01 paired t-test, n = 16 and 17; *p < 0.05, Mann-Whitney U test, n = 17).

L-NAME administration attenuated the recovery-induced increase in choroidal IL-6 transcription, suggesting that NO is involved in the regulation of choroidal IL-6 mRNA transcription. Therefore, as a second approach to evaluate NO on choroidal IL-6 transcription, we tested the effect an NO donor on IL-6 gene transcription in isolated chicken choroids (**Figure 7**). Treatment of choroids with PAPA-NONOate, an NO donor with a half-life of 15 min at 37 ° C, led to a concentration dependent increase in IL-6 mRNA that reached a 5-fold increase at 1.5 mM (**Figure 7A**). Protein expression of IL-6 was also significantly increased in isolated choroids following incubation with PAPA-NONOate, compared to that of choroids incubated in culture medium alone (↑ 30.52 %, and ↑131.77% for control vs 3 mM PAPA-NONOate and control vs 5 mM PAPA-NONOate; p < 0.05, p< 0.0.01, respectively, ANOVA with Bonferonni’s correction), (**Figure 7B**).

**Figure 7.**
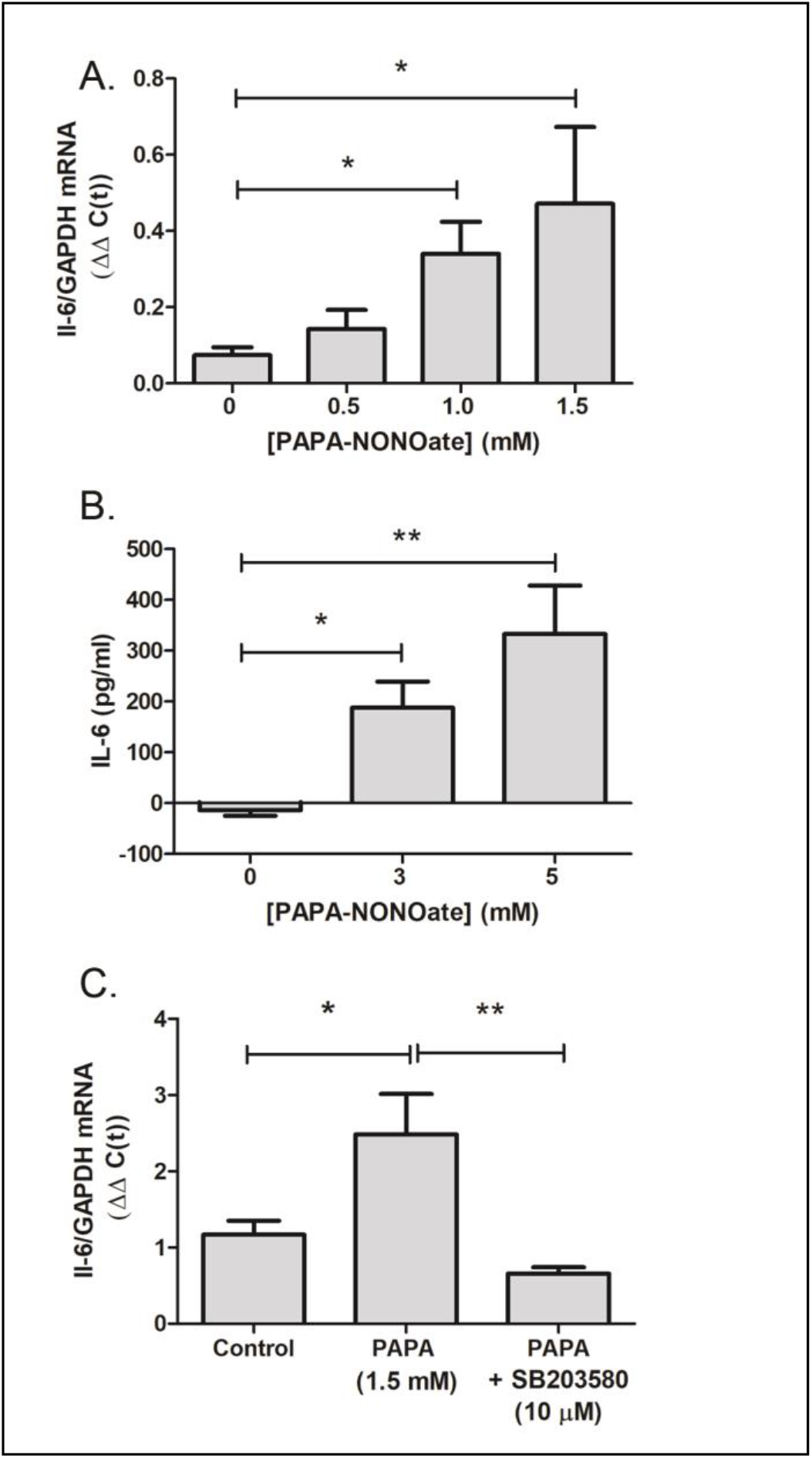
The NO donor, PAPA-NONOate, stimulates choroidal IL-6 production. Choroids were isolated from normal chicken eyes and were incubated with the indicated concentrations of PAPA-NONOate for 24 hrs. **(A)** IL-6 gene expression was significantly increased in choroids following incubation in 1.0 and 1.5 mM PAPA-NONOate (*p < 0.05, ANOVA with Bonferonni’s correction, n = 4 - 5 choroids in each group). **(B)** IL-6 protein concentrations were significantly increased in choroid culture supernatants following incubation with 3 – 5 mM PAPA-NONOate (**p < 0.01, *p <0.05, ANOVA with Bonferonni’s correction, n = 3 - 5 choroids in each group). **(C)** Incubation of chicken choroids with PAPA-NONOate (1.5 mM) together with the p38 MAPK inhibitor SB203580 (10 μM) abolished the PAPA-NONOate-induced increase in IL-6 mRNA (**p < 0.01, * p< 0.05, Student’s t-test, n = 10 choroids in each group).

As NO has been shown to activate members of the MAPK pathway in a cGMP-independent manner, and given that the p38 pathway plays essential roles in the production of IL-6 and other proinflammatory cytokines (IL-1β, TNF-α and IL-6) (Guan, Buckman et al., 1998), we sought to determine whether p38 MAPK activation contributes to the NO mediated stimulation of choroidal IL-6 transcription. Treatment of isolated choroids with the p38 specific inhibitor, SB203580, and PAPA-NONOate abolished the NO-induced increase in IL-6 mRNA, suggesting that the NO-stimulated IL-6 transcription is mediated through activation of MAPK signaling pathways (**Figure 7C**).

### Atropine Stimulates Choroidal Il6 Transcription

Atropine has been shown to be clinically effective at reducing myopia progression in clinical trials (Upadhyay & Beuerman, 2020) and in avian and mammalian animal models of myopia, although the mechanism of action is poorly understood (McBrien, Moghaddam et al., 1993, Whatham, Lunn et al., 2019). Therefore, we examined the effect of atropine on choroidal IL-6 transcription in chicks undergoing form deprivation myopia (**Figure 8**). Intravitreally delivered atropine (240 nmol/eye) significantly increased choroidal IL-6 mRNA in form deprived eyes as compared with vehicle-treated form deprived eyes (↑81%; p <0.05), when measured 6 hrs following atropine administration (**Figure 8A**). Interestingly, application of atropine (0.1%) directly to isolated chick choroids stimulated IL-6 gene expression (↑103%, p < 0.01, Student’s t-test) (**Figure 8B**).

**Figure 8.**
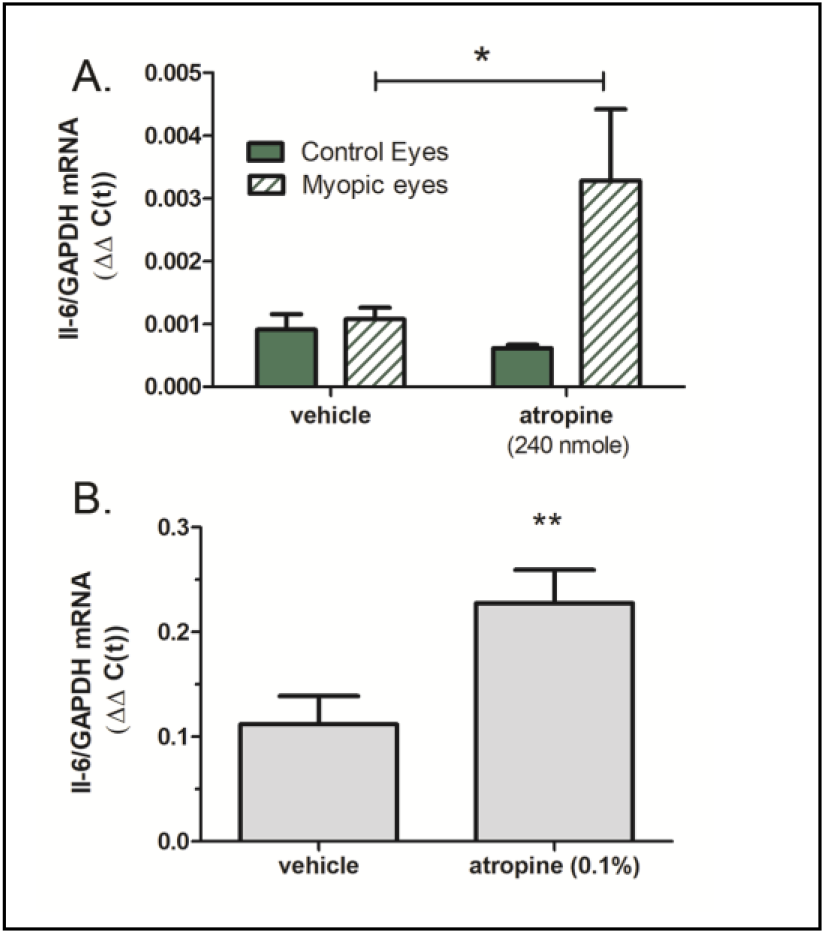
Atropine stimulates choroidal IL-6 gene expression. **(A)** Intravitreal injection of atropine (240 nmole/eye) into chick eyes following 14 days for form deprivation (Myopic eyes) increased IL-6 mRNA levels compared to myopic eyes receiving vehicle only (PBS) (*p < 0.05, Mann-Whitney U test, n = 18). **(B)** Incubation of chicken choroids in organ culture with 0.1% atropine for 24 hrs significantly increased choroidal IL-6 gene expression. (**p < 0.01, Student’s t-test, n = 16).

## Discussion

These studies document, for the first time, that the potent inflammatory cytokine IL-6 is expressed in and released by the choroid during the recovery from form deprivation myopia. Choroidal upregulation of IL-6 is rapid; significant increases in gene expression were observed after only 45 minutes of unrestricted vision. The visual stimulus for choroidal IL-6 gene expression is myopic defocus, as a similar rise in IL-6 is observed after 6 hrs of positive (+15D) lens wear. Finally, nitric oxide appears to directly or indirectly upregulate choroidal IL-6 gene expression.

We and others have previously shown that changes in the visual environment can cause rapid changes in ocular growth and refraction as evidenced by changes in scleral proteoglycan synthesis, choroidal retinoic acid synthesis, and choroidal thickness [*reviewed in* (Troilo et al., 2019)]. The identification of a chemical mediator of these visually induced changes in ocular growth has been elusive. IL-6 is a pleiotropic cytokine, synthesized by a variety of cell types, that plays key roles in immune responses, inflammatory reactions, as well as the growth and differentiation of many cell types (Kishimoto, 2006). For these reasons, choroidally derived IL-6 is an attractive candidate as a potential mediator in the retina-to-sclera signaling cascade. Here we demonstrate that myopic defocus, as a result of prior form vision deprivation, or due to application of +15D lenses, stimulates choroidal IL-6 gene expression. In contrast, induction of hyperopic defocus through the application of −15D lenses resulted in a slight decrease in choroidal IL-6 gene expression, that did not reach statistical significance in the present study. Choroidal IL-6 gene expression was not significantly affected by rearing chicks in varying light intensities (5 – 3150 lux), or rearing in constant red or blue LED light (58 and 111 lux, respectively), but was significantly lower in choroids from chicks reared in constant darkness for 6 hrs, compared with birds reared in any other light conditions examined. Protein levels of IL-6 were also significantly increased in choroids of recovering eyes following 6 hrs of recovery, but were not significantly different following 24 hrs of recovery. We suspect that IL-6 protein is released from choroids and either enters the circulation or adjacent ocular tissues shortly after its synthesis as has been described for IL-6 in skeletal muscle (Steensberg, van Hall et al., 2000).

The observed elevation in choroidal IL-6 gene and protein expression in response to recovery/myopic defocus does not appear to be due to a general inflammatory response, as other pro-inflammatory cytokines, IFN-γ and TNF-α were unaffected by visual manipulations. IL-1B was significantly elevated following 6 hours of recovery, but was unaffected at 1.5 and 3 hrs of recovery, suggesting that changes in IL-1B gene expression are downstream to that of IL-6.

The finding that myopic defocus significantly upregulated choroidal IL-6 transcription, but hyperopic defocus had only a modest effect at downregulating choroidal IL-6, may be related to the observation that myopic defocus is a more potent stimulus than hyperopic defocus at causing changes in choroid thickness, with regard to the duration of lens wear needed to elicit changes in choroidal thickness, as well as the duration of the response to lens wear (Zhu, Park et al., 2005). If IL-6 is involved in mediating changes in choroidal thickness, we would predict that myopic defocus-induced increases in choroidal IL-6 gene and protein expression would induce changes in choroidal thickness that would outweigh and outlast the minor reduction in IL-6 expression induced by brief periods of hyperopic defocus.

If choroidal IL-6 gene expression is causally related to ocular changes associated with recovery from myopia, then agents known to block recovery should also block choroidal IL-6 upregulation. To test this, we evaluated choroidal IL-6 gene expression in recovering eyes following intravitreal application of the nitric oxide synthase inhibitor, L-NAME. L-NAME has previously been shown to prevent choroidal thickening and disinhibit scleral proteoglycan synthesis during recovery from induced myopia and in response to positive lens wear (Nickla, Wilken et al., 2006). We found that intravitreal treatment with L-NAME significantly attenuated the recovery-induced increase in choroidal IL-6 gene expression, indicating that nitric oxide either directly or indirectly regulates choroidal IL-6 gene expression. Treatment of isolated choroids with the NO donor, PAPA-NONOate stimulated IL-6 gene and protein expression, confirming that choroidal IL-6 is upregulated by NO. NO has been reported to stimulate IL-6 production in skeletal myocytes (Makris, Sotzios et al., 2010), human blood mononuclear cells (Siednienko, Nowak et al., 2011) and kidney epithelial cells (Demirel, Vumma et al., 2012) by activating ERK1/2 and p38 MAPK-signaling pathways. We also found that p38 MAPK signaling was crucial for the PAPA-NONOate-mediated activation of IL-6 in chick choroids as incubation with the p38 MAPK inhibitor, SB203580, abolished PAPA-NONOate-stimulation of IL-6 gene expression.

In the present study, since L-NAME was delivered intravitreally and NO was applied directly to the choroid, it is unclear as to cellular source of NO generated in response to myopic defocus that is responsible for IL-6 upregulation. NO and nNOS (NOS1) have been detected in virtually all retinal neurons, in the RPE, and in several cell types in the chick choroid (Fischer & Stell, 1999, WD, 2000). It is unlikely that choroidally-derived NO is responsible for IL-6 upregulation in the choroid, as application of the NO substrate, L-arginine (which is catalyzed to L-citrulline and NO by the NOS enzymes) had no significant effect on choroidal IL-6 synthesis (**Supplementary Figure S1**). We speculate that NO, released from the retina or RPE diffuses to the choroid to stimulate IL-6 synthesis. It is also unclear as to the cellular source of choroidal IL-6 in response to myopic defocus. Our immunohistochemical evaluation of IL-6 protein distribution indicated that IL-6 was present as discrete puncta in the RPE, choroidal vascular endothelial cells and extravascular stromal cells. Any of these cells, as well as myeloid and lymphoid cells, could be the source of visually induced IL-6. However, IL-6-positive choroidal cells identified in the present study may indicate internalization of IL-6 following synthesis and secretion by neighboring cells via a paracrine signaling mechanism.

Atropine has been shown to prevent experimentally-induced myopia in chicks and inhibits myopia development in some children when applied topically (Chia, Chua et al., 2012). We therefore evaluated choroidal IL-6 gene expression following *in vivo* and *in vitro* treatment with atropine. We found that a single intravitreal injection of atropine to chicks undergoing form deprivation-induced myopia stimulated choroidal IL-6 gene expression. Moreover, incubation of isolated choroids with 0.1% atropine also stimulated choroidal IL-6 gene expression, suggesting that atropine acts directly on the choroid to stimulate IL-6 gene expression.

## Conclusions

In the present study, we report that myopic defocus, either in eyes recovering from induced myopia, or in eyes treated with +15D spectacle lenses, stimulates IL-6 mRNA and protein synthesis in the chick choroid. The ramifications of increased choroidal IL-6 synthesis are unclear. In the context of ocular growth control, it appears that choroidal IL-6 is associated with a slowing of eye growth, as it is upregulated in recovering eyes (when eyes are decelerating their rate of elongation) and in myopic eyes treated with atropine, an agent known to inhibit vitreous chamber elongation and myopia. Moreover, IL-6 mRNA is downregulated in recovering eyes treated with L-NAME, a compound known to inhibit recovery and increase scleral proteoglycan synthesis and ocular elongation. Treatment of isolated sclera with IL-6 had no effect on scleral proteoglycan synthesis (**Supplementary Figure S2**), indicating that additional downstream mediators, most likely derived from the choroid, are responsible for regulating the scleral changes associated with recovery.

On the other hand, studies have shown that IL-6 has a major role in the pathology of uveitis, glaucoma, retinal vein occlusion, macula edema and diabetic retinopathy (Zahir-Jouzdani, Atyabi et al., 2017). IL-6 induces ocular inflammatory responses often leading to the breakdown of the blood ocular barrier, angiogenesis, increased vascular permeability and choroidal neovascularization. Based on the results of the present study, it is possible that myopic defocus in humans, as a result of uncorrected myopia, may cause elevated choroidal IL-6 which could predispose individuals to one or more of the above serious ocular complications.

The identification of small molecule or biological approaches to manipulate choroidal IL-6 concentrations will elucidate the role of choroidal IL-6 in postnatal ocular growth, as well as in a variety of ocular conditions.

## Materials and methods

### Ethics and Animals

Animals were managed in accordance with the ARVO Statement for the Use of Animals in Ophthalmic and Vision Research, with the Animal Welfare Act, and with the National Institutes of Health Guidelines. All procedures were approved by the Institutional Animal Care and Use Committee of the University of Oklahoma Health Sciences Center. White Leghorn male chicks (*Gallus gallus*) were obtained as 2-day-old hatchlings from Ideal Breeding Poultry Farms (Cameron, TX). Chicks were housed in temperature-controlled brooders with a 12-hour light/dark cycle and were given food and water ad libitum. At the end of experiments, chicks were euthanized by overdose of isoflurane inhalant anesthetic (IsoThesia; Vetus Animal Health, Rockville Center, NY), followed by decapitation.

### Visual Manipulations

Form deprivation myopia (FDM) was induced in 3 to 4 day-old chicks by applying translucent plastic goggles to one eye, as previously described (Rada, Thoft et al., 1991). The contralateral eyes (left eyes) of all chicks remained untreated and served as controls. Chicks were checked daily for the condition of the goggles. Goggles remained in place for 10 days, after which time the goggles were removed and chicks were allowed to experience unrestricted vision (recover) for up to 4 days. When multiple time points were assessed in one experiment, chicks were randomly assigned to groups for each time point.

Lens induced myopia and hyperopia were induced via the application of +15 and −15D lenses to 3 to 4 day old chicks. Lenses were fashioned from PMMA hard contact lenses [12 mm diameter, 8 mm base curve, Conforma Labs, Inc (Norfolk, VA)] that were mounted onto nylon washers for support using optical adhesive (Norland Products, Inc., Cranbury, NJ). A velcro ring was glued to the back of the nylon washer for mounting around the chick’s right eye using cyanoacrylate adhesive. Lenses remained in place for up to 24 hrs.

For light intensity experiments, cages (24”×24”×16”, L × W × H, respectively) were fitted with Multicolor (RGB) and White LED strip lights at the top surface of the cage and light intensity was controlled using a wireless RF remote (Super Bright LEDs, Inc., St. Louis, MO). Light intensity (5 lux – 3150 lux) was measured using a light meter (Datalogger Model 401036, Extech Instruments, Nashua, NH) at a distance of 8 cm from the bottom of the cage (approximate eye-level of chicks). Chicks were randomly assigned to white, red or blue light and housed in LED cages for 6 hrs (9:30am – 3:30 pm). A separate group of chicks was kept in complete darkness for 6 hrs.

### Intravitreal injections

Injections were delivered using a NanoFil-100 syringe with a 26G needle (World Precision Instruments, Sarasota, FL) under isofluorane (0.8% in O_2;_ IsoThesia; Vetus Animal Health, Rockville Center, NY) inhalation anesthesia at a flow rate of 0.4 liters/minute using an Isoflurane Anesthesia machine for veterinary use only (Ohmeda Anesthesia Service and Equipment, Inc., Atlanta, GA). Following removal of the occluder, the sclera was exposed by retracting the eyelids with a handmade ocular speculum and injections were delivered through the sclera at the superior margin of the globe, just outside of the scleral ossicles, after cleaning the eye lids and surround area with 70% alcohol. Injections consisted of L-NAME (Sigma Chemical Co., St. Louis, MO) (a 30 μl injection containing 16.2 μmole of L-NAME in 0.9% saline), 30 μl of 0.9% saline (vehicle for L-NAME) (Nickla & Wildsoet, 2004), atropine sulfate (Sigma Chemical Co.) (a 20 μl injection containing 240 nmoles of atropine sulfate in phosphate buffered saline, PBS), and 20 μl of PBS (vehicle for atropine sulfate)(Carr & Stell, 2016). The needle remained in place for 15 seconds before slowly withdrawing it from the eye and an ophthalmic antibiotic ointment (Vetropolycin, Pharmaderm, Melvill, New York) was applied to the eye. In some cases, the occluders were replaced prior to prior to awakening from the anesthesia.

### Tissue Preparation

Chicks were euthanized by an overdose of isoflurane inhalant anesthetic (IsoThesia; Vetus Animal Health) following 10 days of form deprivation (day 0 recovery), after various time points of recovery, following lens wear, or light exposure. Eyes were enucleated and cut along the equator to separate the anterior segment and posterior eye cup. Anterior tissues were discarded, and the vitreous body was removed from the posterior eye cups. An 8 mm punch was taken from the posterior pole of the chick eye using a dermal biopsy punch (Miltex Inc., York, PA). Punches were located nasal to the exit of the optic nerve, with care to exclude the optic nerve and pecten oculi. With the aid of a dissecting microscope, the retina and majority of RPE were removed from the underlying choroid and sclera with a drop of phosphate buffered saline (PBS; 3 mM dibasic sodium phosphate, 1.5 mM monobasic sodium phosphate, 150 mM NaCl, pH 7.2) and gentle brushing. For microarray, Taqman real time PCR, and ELISA assays, choroids were separated from the sclera using a small spatula, placed in 2 ml screw cap tubes, and snap frozen in liquid nitrogen and stored at −80°C. For immunolabelling experiments, choroids with sclera still attached were placed into a 48-well flat bottom plate (Corning Inc., Corning, NY). A small amount of RPE was left on the choroids to discriminate between the RPE and scleral side of the tissue. The tissues were then fixed with 4% paraformaldehyde (stock solution freshly prepared) in PBS O/N at 4°C.

### Immunolabelling of Chick Choroids

Punches (5 mm) containing retina, RPE choroid, and sclera were obtained from the posterior poles of control and recovering chick eyes, fixed in neutral-buffered formalin, and embedded in paraffin, and sections were obtained. Tissue sections of posterior ocular tissues were deparaffinized through a graded series of xylenes and ethanol and rinsed in PBS, and then incubated for 30 min at RT in incubation buffer that consisted of 2% BSA (Sigma Chemical Co.) and 0.2% Triton X-100 in PBS. Sections were incubated overnight at 4 °C with rabbit anti-chick IL-6 (Bio-Rad Laboratories, Inc., Hercules, CA) diluted 1:20 in incubation buffer. For negative controls, tissue sections were incubated in 25 μg/ml nonimmune rabbit immunoglobulin (Sigma Chemical Co.) instead of the IL-6 antibody. Additional pre-absorption controls were performed in which the anti-IL-6 antibody was incubated overnight at 4 °C with a 10 fold molar excess of recombinant chicken IL-6 (1.67 μM; Bio-Rad Laboratories, Inc.) before immunolabeling fixed sections of chick ocular tissues. Following overnight incubation with the primary antibody, sections were rinsed in PBS, and incubated for 30 min at RT in 5 μg/ml of goat anti-rabbit AlexaFluor 488 (ThermoFisher Scientific, Richardson, TX). Sections were rinsed in PBS and then incubated for 10 s at RT with 0.0005% DAPI nuclear stain, followed by a final rinse in PBS. Coverslips were mounted onto the slides with Prolong Gold Antifade reagent containing DAPI (ThermoFisher Scientific), and the immunolabeled sections were examined under an Olympus Fluoview 1000 laser-scanning confocal microscope (Center Valley, PA).

### Microarray Analyses

Choroids were isolated from 10 normal chick eyes (n = 5 chicks) and from control and treated chicks eyes following 6 hrs of recovery from 10 days of prior form deprivation-induced myopia (n = 5 chicks) and kept at −80° C until processed. Choroids were shipped on dry ice to the Microarray Core Facility at the University of Tulsa (Tulsa, Oklahoma). When processing began the samples were moved to a container of liquid nitrogen. The choroids were pulverized using a frozen 1.5mL disposable pestle. Immediately following pulverization the samples were immersed in 300μL of Ambion TriReagent (Applied Biosystems, Foster City, CA) solution and homogenized for 90 seconds with a Pellet Mixer. An additional 700μL of TriReagent was pipetted into the sample after homogenization. Incubation of samples occurred for 5 min using a 1.5 mL microfuge tube shaker at room temperature. The samples were then transferred to pre-spun Phase Lock Gel Heavy 2mL Gel tubes (5 Prime Inc., Gaithersburg, MD). 200μL of chloroform was added to each sample, inverted 12 times, and incubated at room temperature for 5 min. The samples were then spun at 2° C for 20 min. 500μL supernatant was poured into 2mL round bottom tubes. These tubes were placed into the Qiagen Qiacube robotic workstation and cleaned using the RNeasy Lipid Tissue Mini Kit (Qiagen, Redwood City, CA). The samples were eluted in 50μL of molecular biology water. The samples were also split into two 25μL aliquots to ensure sample safety.

Following RNA isolation, the samples were quantified using a NanoDrop 1000 spectrophotometer (ThermoFisher Scientific). The initial average sample concentration ranged from 20 – 88 ng/μL. The initial RNA 260/280 ratios were between 1.8 and 2.0 with the 260/230 ratios between 0.08 and 1.9. Precipitation of one aliquot of RNA was performed to increase the sample concentration and purity. This procedure was performed by addition of 2.5 volumes of ice cold 100% EtOH, 1/10 of 3M ammonium acetate, and 1μL of glycogen at 5 ng/μL. The samples were incubated at −20° C overnight. The samples were spun at 4° C for 30 min to pellet the RNA. The supernatant was removed and the pellet was washed with ice cold 80% EtOH to remove remaining salt. The EtOH was aspirated off and the pellet was dried at room temperature for 5 min. Molecular biology water was used to re-suspend the RNA pellet. The amount of water used was calculated to bring the sample concentration to between 58 – 133 ng/μL. After precipitation the 260/280 ratios are between 2.0 and 2.1 and the 260/230 ratios are between 1.8 and 2.1. 150 ng of each sample was and processed with the Affymetrix 3’ IVT Express Kit (ThermoFisher Scientific).

### TaqMan Quantitative PCR (RT-Quantitative PCR)

Choroids were isolated from individual pairs of control and treated eyes and snap frozen in liquid nitrogen. Total RNA was isolated using TRIzol reagent (ThermoFisher Scientific) followed by DNase treatment (DNA-free, Applied Biosystems) as described previously (Summers, Harper et al., 2016). RNA concentration and purity were determined via the optical density ratio of 260/280 using a Nanodrop ND-1000 spectrophotometer and stored at −80 °C until use. cDNA was generated from DNase-treated RNA using a High Capacity RNA to cDNA kit. Real time PCR was carried out using a Bio-Rad CFX 96. 20-μl reactions were set up containing 10 μl of TaqMan 2× Universal Master Mix (Applied Biosystems), 1 μl 20× 6-carboxyfluorescein (FAM)-labeled Assay Mix (Applied Biosystems), and 9 μl of cDNA. Each sample was set up in duplicate with specific primers and probed for chicken IL-6 (assay ID number Gg03337980_m1), chicken interferon γ (INFG, assay ID number Gg03348618_m1), chicken IL-1β (IL1β, assay ID number Gg03347154_g1), chicken TNF-α (LITAF, assay ID number Gg03364359_m1) and the reference gene chicken GAPDH (assay ID number Gg03346982_m1) (Thermofisher Scientific). The PCR cycle parameters were an initial denaturing step at 95 °C for 10 min followed by 45 cycles of 95 °C for 15 s and 60 °C for 1 min. Normalized gene expression was determined by the ΔΔ*c*(*t*) method (Livak & Schmittgen, 2001) using Bio-Rad CFX Manager™ version 3.1 and reported values represent the average of duplicate samples.

### IL-6 protein measurements

Punches (8 mm) of chick choroids were rinsed in ice-cold PBS (0.01 M, pH 7.2) and homogenized in 300 μl of PBS on ice (Omni Tip, Omni International, Kennesaw, GA). The resulting suspension was sonicated with an ultrasonic homogenizer (Pulse 150, Benchmark Scientific, Edison, NJ) and subjected to two freeze-thaw cycles to further break the cell membranes. Homogenates were then centrifugated for 5 minutes at 5000 × g. Following centrifugation, pellets were discarded and the supernatants stored at ≤−20°C. IL-6 was measured on duplicate samples using a commercially available chicken IL-6 ELISA kit (Aviva Systems Biology, Corp., San Diego, CA) according to the manufacturer’s instructions. Protein concentrations in choroidal lysates were determined on duplicate samples by Bradford assay. Reported values represent the average of duplicate samples.

### Organ Culture

Choroids were isolated from eyes from adult chicken heads, (Animal Technologies, Inc., Tyler, Texas) as described above and placed in 48-well plates containing 300 μl culture medium [1:1 mixture of Dulbecco’s Modified Eagle’s Medium and Ham’s F12 containing streptomycin (0.1 mg/ml), penicillin (100 units/ml) and gentamicin (50 ug/ml)] in the presence of the NO donor, PAPA-NONOate (0.5 – 5 mM in culture medium; Cayman Chemical, Ann Arbor, MI), the p38 MAPK inhibitor, SB203580 (10 μM; Sigma), atropine sulfate (0.1%; Sigma) or culture medium alone in a humidified incubator with 5% CO_2_, overnight at 37 °C. Following incubation, choroids were snap frozen and RNA isolated for TaqMan real time PCR assays, and medium harvested and frozen for IL-6 ELISA assays.

### Scleral Sulfated Glycosaminoglycan Synthesis

The posterior hemispheres of eyes of FD chicks (= 0 days of recovery), or from eyes from chicks recovering from FD myopia for 1 – 20 days and contralateral controls were obtained and one 5 mm tissue punch was excised from the posterior sclera of control and treated eyes using a dermal punch (Miltex Instrument Co.). All retina, RPE, choroid, vitreous, pectin, and muscle were gently cleaned from each sclera punch. Scleral punches were initially placed into wells of a 96 well culture plate with 50 μl of N2 medium [Ham’s F-12/DMEM containing 1x N2 supplement (Stem Cell Technologies, Vancouver, BC) until all sclera were obtained. Scleral punches were then transferred to N2 medium containing ^35^SO_4_ (100 μCi/ml; New England Nuclear, MA) and incubated for 3 hr. at 37°C. Radiolabelled scleral punches were digested with proteinase K (protease type XXVIII, Sigma Chemical Co.), (0.05% w/v in 10 mM EDTA, 0.1 M sodium phosphate, pH 6.5) overnight at 60°C. ^35^SO_4_ –labeled glycosaminoglycans (GAGs) were precipitated by the addition of 0.5% cetylpyridinum chloride (CPC) in 0.002 M Na^2^S0_4_ in the presence of unlabeled carrier chondroitin sulfate (1mg/ml in dH_2_O). The samples were incubated for 30 min at 37°C and precipitated GAGs were collected on Whatman filters (GF/F) using a Millipore 12-port sampling manifold as previously described (Rada, McFarland et al., 1992). Radioactivity was measured directly on the filters by liquid scintillation counting.

### Statistics

Sample sizes were calculated using G*Power 3.1.9.2 using two tailed tests with an α = 0.05, and an effect size determined by group means and standard deviations previously published by this lab and others (Rada, Thoft et al., 1991, Wallman & Adams, 1987. All experiments were repeated at least one time, and sample sizes and results reported reflect the cumulative data for all trials of each experiment. Parametric analyses between groups were made using paired or unpaired Student’s t-tests, and a one-way ANOVA followed by a Bonferroni correction for multiple comparisons. Non-parametric analyses between groups were made using the Mann-Whitney U test, or the Kruskal-Wallis test for multiple comparisons (GraphPad Prism 5, La Jolla, CA). Results were considered significant with p-value ≤ 0.05.

## Acknowledgements

This work was supported by NIH grant R01EY09391 (JAS) and by NIGMS COBRE Grant P30GM122744 (Ma, J-X., PI). The authors would like to thank Dr. Frederick (Kris) Miller (Department of Cell Biology, University of Oklahoma Health Science Center) and Dr. Randle Gallucci (Department of Pharmaceutical Sciences, University of Oklahoma Health Science Center) for their helpful discussions and suggestions.

## Supplementary Figures

**Figure S1.**
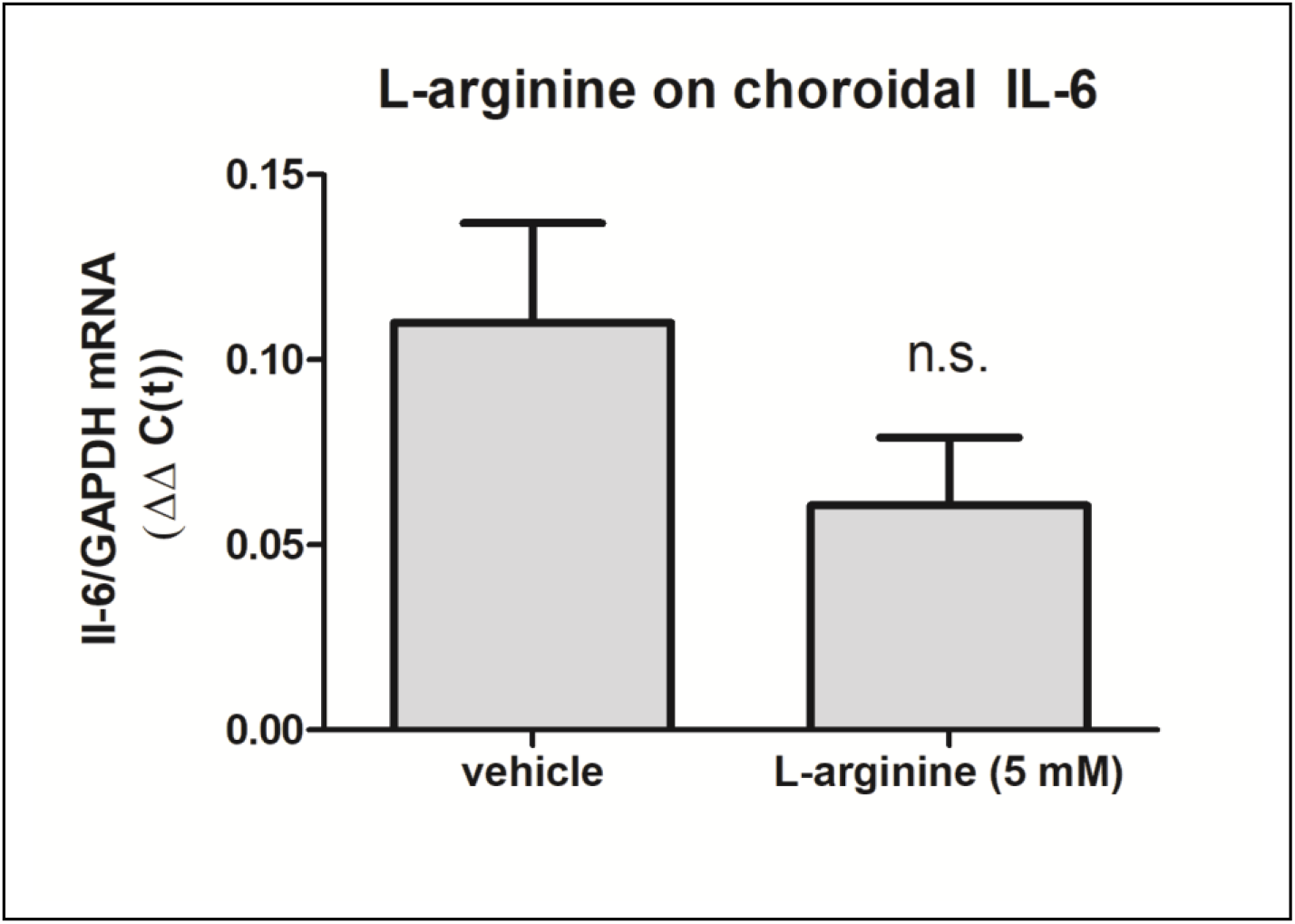
L-arginine does not stimulate choroidal IL-6 gene expression. Incubation of chicken choroids in organ culture with L-arginine for 24 hrs had no significant effect on choroidal IL-6 gene expression. (p = 0.1401, Student’s t-test, n = 16 choroids in each group).

**Figure S2.**
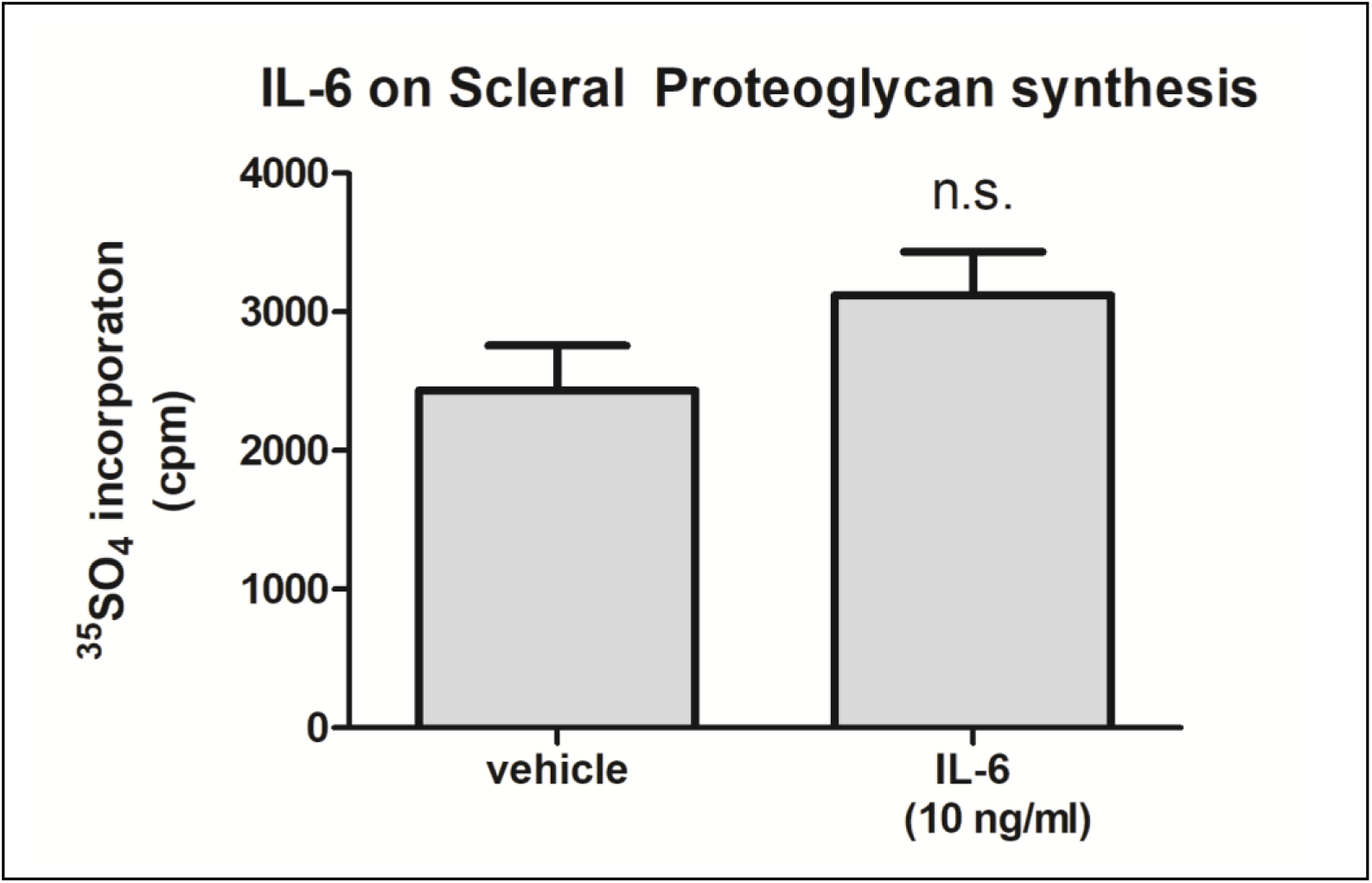
IL-6 has no direct effect on scleral proteoglycan synthesis. Incubation of chicken sclera in organ culture with recombinant chicken IL-6 (10 ng/ml) for 24 hrs had no significant effect on scleral proteoglycan synthesis. (p = 0.1439, Student’s t-test, n = 11 sclera in each group).

